# Enteropancreatic neurons drive the glucoregulatory response to ingested lipid

**DOI:** 10.1101/2025.05.09.652620

**Authors:** Anna G. Roberts, Leah Meyer, Jieruo Liu, Mariana Norton, Phyllis Phuah, Aldara Martin Alonso, Georgina K.C. Dowsett, Sijing Cheng, Cecilia Dunsterville, Pei-En Chung, Yuxuan Tao, Tobias Smitherman-Cairns, Alexandra B. Deutsch, Abhisekh Chatterjee, Brian Y.H. Lam, Aylin C. Hanyaloglu, Ben Jones, Giles S.H. Yeo, Victoria Salem, Kevin G. Murphy

## Abstract

Enteropancreatic neurons project from the small intestinal wall to the pancreas. Though well positioned to mediate the effects of ingested nutrients on pancreatic function, the metabolic role of these neurons is unclear. Diets rich in olive oil promote weight loss and improve remission rates in patients with T2DM. Here, we show that olive oil improves acute glucose tolerance by stimulating insulin release via neurotensin receptor type 1 (NTSR1)-expressing enteropancreatic neurons. These neurons are necessary for the effects of olive and neurotensin on glucose tolerance, and their activation is sufficient to improve glucose tolerance. These findings suggest a mechanism by which dietary olive oil regulates blood glucose levels and present a novel functional role for enteropancreatic neurons in regulating glucose homeostasis.

## Introduction

The Mediterranean Diet, rich in vegetables, nuts, fruit, whole grains and olive oil, reduces metabolic syndrome indicators, including reduced risk of cardiovascular disease, alongside improvements in weight management and T2DM remission^1–3^. Evidence suggests that olive oil may be the crucial component for improved glucoregulation. A Mediterranean Diet supplemented with additional olive oil further reduced the incidence of T2DM and chronic adherence to olive-oil enriched diets lowers blood glucose levels^4,5^.

Oleic acid is the main monounsaturated fatty acid (MUFA) within the Mediterranean Diet^6^ and is present at high levels in olive oil. Previous work has shown both olive oil- and oleic acid induce secretion of the gut hormone neurotensin from enteroendocrine cell models and in vivo in a range of species^7–10^. Neurotensin is a 13-amino acid peptide synthesized in the gastrointestinal tract and the central nervous system. Most known biological effects of neurotensin are mediated by the neurotensin receptor type 1 (NTSR1). Centrally, neurotensin-NTSR1 signalling plays roles in thermoregulation^11^, analgesia^12^, addiction^13^, food intake^14^ and blood pressure regulation^15^, and peripherally has been reported to modulate gastric emptying^16,17^ and lipid absorption^18–20^. However, the role of the neurotensin-NTSR1 system in glucose homeostasis is unclear^21–27^. Here we show that the acute glucoregulatory effects of olive oil are mediated through NTSR1-expressing enteropancreatic neurons that extend directly from the wall of the duodenum to the pancreas and represent a novel pathway by which dietary intake regulates glucose homeostasis. This is the first report of a physiological role for a population of enteropancreatic neurons.

## Results

### Olive oil increases insulin secretion and improves acute glucose tolerance via a NTSR1-mediated pathway

We investigated the effects of olive oil on acute glucose tolerance in mice. Olive oil significantly improved glucose tolerance 15-minutes following intraperitoneal or oral glucose administration in lean mice (Fig.1A, Extended Data Fig.1A). Olive oil also increased circulating insulin levels (Fig.1B). Olive oil is rich in MUFAs which have been suggested to mediate its metabolic effects. A control oil rich in saturated fatty acids rather than MUFAs did not improve glucose tolerance in mice (Extended Data Fig.1B). Of the MUFAs in virgin olive oil, oleic acid comprises approximately 60-80%^28^. We therefore investigated whether oleic acid might be responsible for the glucoregulatory effects of olive oil. Oral co-administration of the lipase inhibitor Orlistat, which prevents the breakdown of fats into fatty acids, blocked the improvement in glucose tolerance seen with olive oil (Fig.1C), and oral administration of oleic acid alone significantly improved glucose tolerance (Fig.1D). Within the gastrointestinal tract, MUFAs are predominantly sensed by the free fatty acid receptor 4 (FFAR4). We found that a specific FFAR4 antagonist also attenuated the glucoregulatory effect of olive oil (Fig.1E). Together these data suggest that the acute effects of olive oil on glucose tolerance are mediated by the sensing of its constituent fatty acids through the FFAR4.

**Figure 1.**
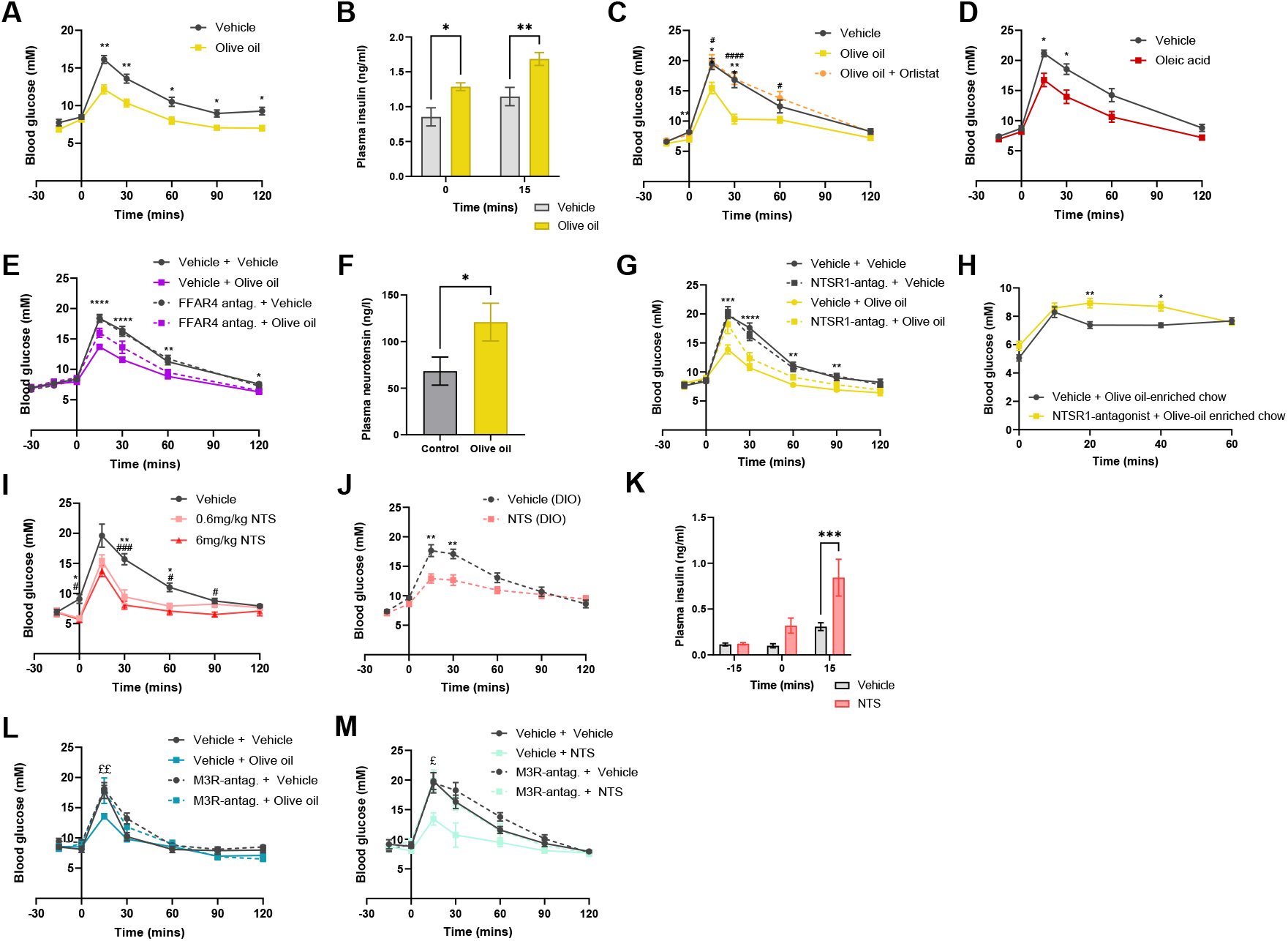
Olive oil improves glucose tolerance via activation of NTSR1. **A).** Intraperitoneal glucose tolerance test (IPGTT) with olive oil or vehicle orally gavaged (OG)-15 minutes prior to glucose load. n=20/group. **B).** Plasma insulin levels measured using Mouse Insulin ELISA following olive oil or vehicle OG administration and intraperitoneal glucose load. n=8-9/group. **C).** IPGTT with olive oil, olive oil and orlistat or vehicle OG-15 minutes prior to glucose load. n=13/group. **D).** IPGTT with oleic acid or vehicle OG-15 minutes prior to glucose load. n=10/group. **E).** IPGTT with subcutaneous administration of FFAR4 antagonist or vehicle 15 minutes prior to olive oil or vehicle OG-15 minutes prior to IP glucose load. n=10/group. **F).** Plasma neurotensin levels measured by ELISA 2 minutes following OG olive oil or vehicle. n=4/group. **G).** IPGTT with 1h pretreatment of an NTSR1-specific antagonist IP followed by olive oil or vehicle OG. n=8/group. **H).** Fast-refeed study with 1h pretreatment of an NTSR1-specific antagonist IP. Mice were refed olive oil-enriched chow at t0 and blood glucose monitored for 1h. n =13/group. **I).** IPGTT with intraperitoneal (IP) neurotensin (0.6mg/kg, 6mg/kg) or vehicle 15 minutes prior to glucose load in lean mice. n=6/group. **J).** IPGTT with IP neurotensin or vehicle 15 minutes prior to glucose load in DIO mice. n=16/group. **K).** Plasma insulin levels measured using Mouse Insulin ELISA following peripheral neurotensin or vehicle administration and intraperitoneal glucose. n=10/group. **L).** IPGTT with 15-minute pretreatment of an M3R-specific or vehicle IP followed by OG olive oil or vehicle n=5/group. **M).** IPGTT with IP 15-minute pretreatment of an M3R-specific antagonist or vehicle followed by IP of either neurotensin or vehicle. n=4/group. All data was analysed with either 1-way or 2-way ANOVA with Tukey’s post-hoc analysis test. * indicates significance between different treatment groups, # indicates significance between olive oil and olive oil + orlistat (Fig. 1C), $ indicates significance between vehicle + olive oil and FFAR-antagonist + olive oil (Fig. 1E) or vehicle + olive oil and NTSR1-antagonist + olive oil (Fig. 1G), £ indicates significance between vehicle + NTS and M3R-antagonist + NTS (Fig. 1L) or vehicle + olive oil and M3R-antag. + olive oil (Fig. 1M). In Fig. 1I #indicates significance between 6mg/kg NTS and vehicle, * indicates significance between 0.6mg/kg NTS and vehicle.

Previous work has shown that both oral and intra-luminal infusions of olive oil and oleic acid increase circulating neurotensin levels^7–10^. We also found that plasma neurotensin levels increased in mice following oral administration of olive oil (Fig.1F). We therefore investigated the role of the NTSR1 in the glucoregulatory response to olive oil. Pretreatment with an NTSR1 antagonist significantly attenuated the glucose-lowering effects of olive oil, suggesting that a significant part of olive oil’s acute glucoregulatory response is controlled by activation of the NTSR1 (Fig 1G). The role of NTSR1 was nutrient specific, as the same NTSR1 antagonist did not block the glucoregulatory effects of an oral protein bolus (Extended Data Fig.1C). Olive oil has also been reported to stimulate the release of the incretin glucagon-like peptide-1 (GLP-1)^29^. To confirm that the majority of the effects of olive oil on glucose tolerance were not mediated by GLP-1, we investigated the effect of a GLP-1 receptor antagonist and conducted glucose tolerance tests in a Pdx1CreERT-GLP-1Rfl/fl mouse model in which the GLP-1R was knocked out of pancreatic beta cells. We observed that olive oil administration still improved glucose tolerance in mice treated with a GLP-1 antagonist, or lacking GLP-1R in beta cells, suggesting GLP-1 signalling is not required for this effect (Extended Data Fig.2G&H).

**Figure 2.**
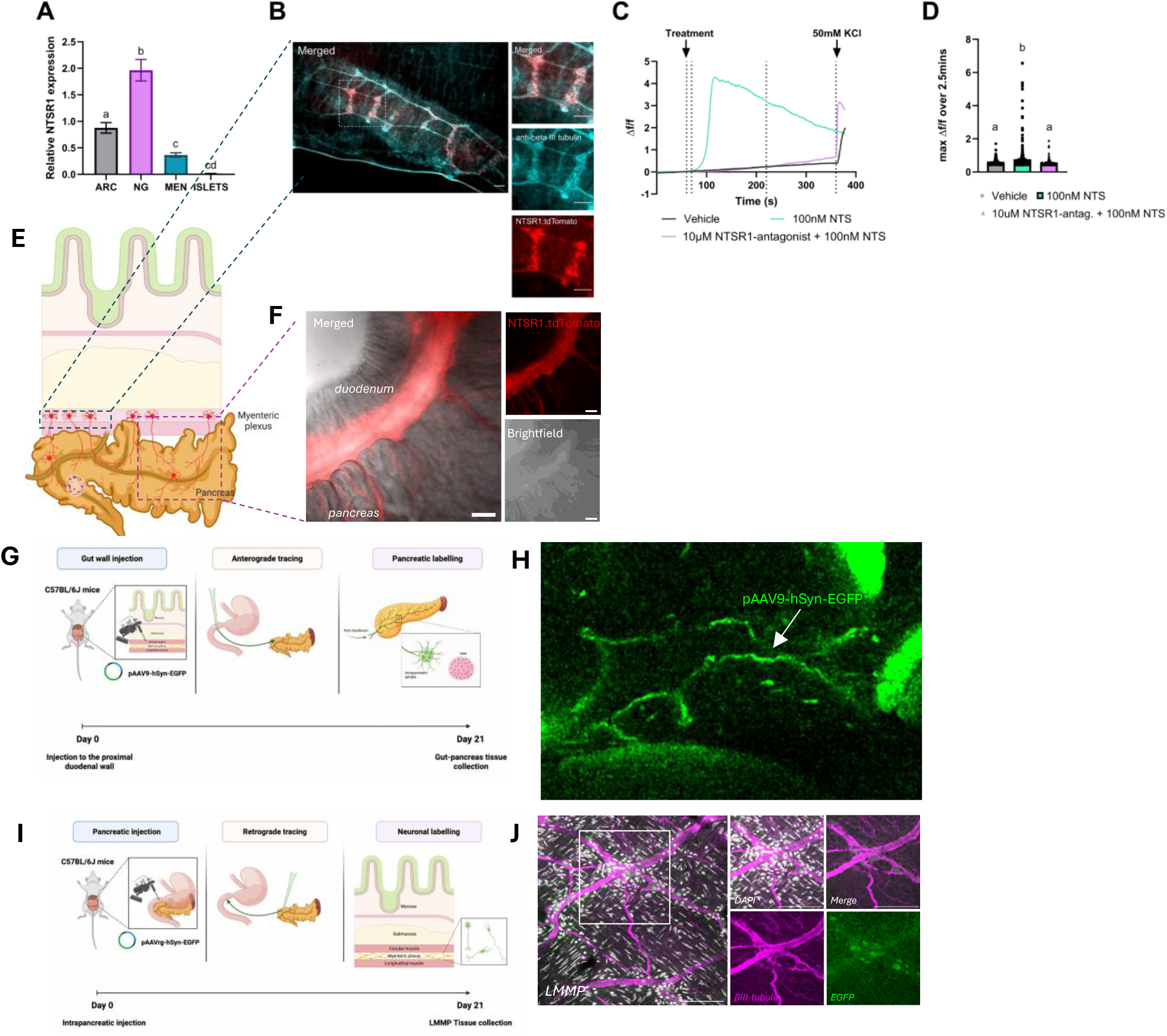
NTSR1 is expressed in enteropancreatic neurons. **A).** qPCR data showing relative NTSR1 expression in the murine arcuate nucleus (ARC), nodose ganglia (NG), primary myenteric enteric neurons (MENs) and isolated islets (ISLETS). n=5-6. **B).** NTSR1 and beta-III tubulin expression in LMMP of the proximal duodenum. **C).** Representative traces of Fura-4 AM-loaded myenteric enteric neurons (MENs) treated with either vehicle (HKRB), 100nM neurotensin or 10µM SR48692 + 100nM neurotensin. **D).** Maximum change in fluorescence measured in the 2.5 minutes following treatment stimulation. 3x technical replicates over 3x biological replicates. **E).** Schematic of NTSR1cre:tdTomato enteropancreatic connection. **F).**Expression of tdTomato in the NTSR1-cre:tdTomato mouse in the enteropancreatic connection at the proximal duodenum in cleared tissue. **G).** Schematic depicting microinjection of anterograde tracer (pAAV9-hSyn-EGFP) into the gut wall. **H).** Expression of eGFP-expressing enteropancreatic neurons advancing into the pancreas. **I).** Schematic depicting microinjection of retrograde tracer (pAAVrg-hSyn-EGFP) into the pancreas. **J).** Expression of eGFP and beta-III tubulin in the LMMP of the proximal duodenum of mice injected into the pancreas with retrograde tracer. All scale bars are 100µm. Change in fluorescence data analysed using 1-way ANOVA with Tukey’s post-hoc analysis.

To assess whether NTSR1 mediates the effects of olive oil in a physiological setting, mice were fed olive-oil enriched chow following an overnight fast. Mice administered an NTSR1 antagonist prior to refeed had higher blood glucose values 1h later, suggesting that olive oil restrains postprandial blood glucose levels in a NTSR1-dependent manner (Fig.1H).

We then investigated the glucoregulatory effects of exogenous neurotensin administration. We found intraperitoneal neurotensin improved acute glucose tolerance in lean mice (Fig.1I). We also found that administration of neurotensin was able to improve glucose tolerance in diet-induced obese (DIO) mice (Fig.1J). Neurotensin administration was also associated with an increase in circulating insulin levels (Fig.1K).

GLP-1R agonists are currently important gut hormone-based pharmacotherapies for T2DM, and there is much interest in combining GLP-1R agonism with other glucoregulatory agents, to increase effectiveness and reduce side effects. Previous work has suggested potential for neurotensin/GLP-1 co-agonism to lower blood glucose levels^26,27^. We similarly confirmed that low-dose neurotensin had an additive effect on glucose tolerance when co-administered with a low-dose of a GLP-1R agonist (Extended Data Fig. 2A).

Neurotensin has been shown to play an important role in controlling lipid absorption in the gut^18–20^. We therefore investigated whether the glucoregulatory effects of peripheral neurotensin were driven by changes in circulating lipid levels that might be sensed at the level of the beta-cell^30^. We found no difference in circulating triglyceride levels in a glucose tolerance test at the time point at which neurotensin or olive oil have glucoregulatory effects (Extended Data Fig.1D&E). Having established that olive oil improves glucose tolerance through a neurotensin-NTSR1-dependent mechanism, we then investigated the specific pathway responsible.

### Neurotensin activates neurons in the myenteric plexus through NTSR1, and NTSR1 is expressed in enteropancreatic neurons

In accord with previous work reporting that NTSR1 mRNA is not expressed in the pancreas itself^31,32^, we found negligible expression in murine pancreatic islets (Fig.2B), suggesting neurotensin does not act directly on the pancreas to improve glucose tolerance. We therefore investigated whether neurotensin might stimulate insulin release through a neural mechanism. The major neural pathway stimulating insulin release acts via the muscarinic-3 acetylcholine receptor (m3R), which is highly expressed on pancreatic beta cells^33,34^. We found that administration of an m3R-antagonist blocked the glucoregulatory effects of both oral olive and peripheral neurotensin (Fig.1L&M). The pancreatic ganglia provide significant direct cholinergic innervation of the pancreatic islets^35^, with the activity of these ganglia modulated by external neuronal populations including vagal neurons extending from the dorsal motor complex (DMC), sympathetic efferent neuronal populations, and neuronal projections that extend directly from the gastrointestinal tract. We were therefore interested to determine whether neurotensin acted centrally, indirectly via afferent vagal signalling, or via enteropancreatic neurons to control glucoregulation.

We confirmed the NTSR1 to be highly expressed in the murine nodose ganglia, arcuate nucleus and myenteric plexus neurons isolated from the proximal duodenum (Fig.2B). Mice in which NTSR1-expressing neurons were ablated in the nodose ganglia, where the cell bodies of afferent vagal neurons reside, showed improved glucose tolerance in response to neurotensin, suggesting that neurotensin does not drive its glucoregulatory effects via direct activation of vagal-NTSR1 (Extended Data Fig.3). Central administration of neurotensin at a dose sufficient to lower body temperature did not improve glucose tolerance, and central blockade of NTSR1 sufficient to block the anorectic effects of peripheral neurotensin did not block the effects of peripheral neurotensin on glucose tolerance. These data suggest peripherally administered neurotensin does not act on the brain to regulate glucose homeostasis (Extended Data Fig.4). In accord with previous reports^36^, we identified NTSR1-expression in the myenteric plexus of the proximal duodenum (Fig.2A&B). It has been reported that NTSR1 immunoreactivity is present in neurons in the pancreas^37^, though NTSR1 mRNA has not been detected^31,32^. We therefore hypothesised that NTSR1 was expressed on enteropancreatic neurons that originate in the wall of the duodenum and that extend to the pancreas to regulate pancreatic function. Enteropancreatic neurons were first reported in both dogs and humans^38^ and later identified in mice^39–41^. However, there has been little study of their physiological role in vivo.

**Figure 3.**
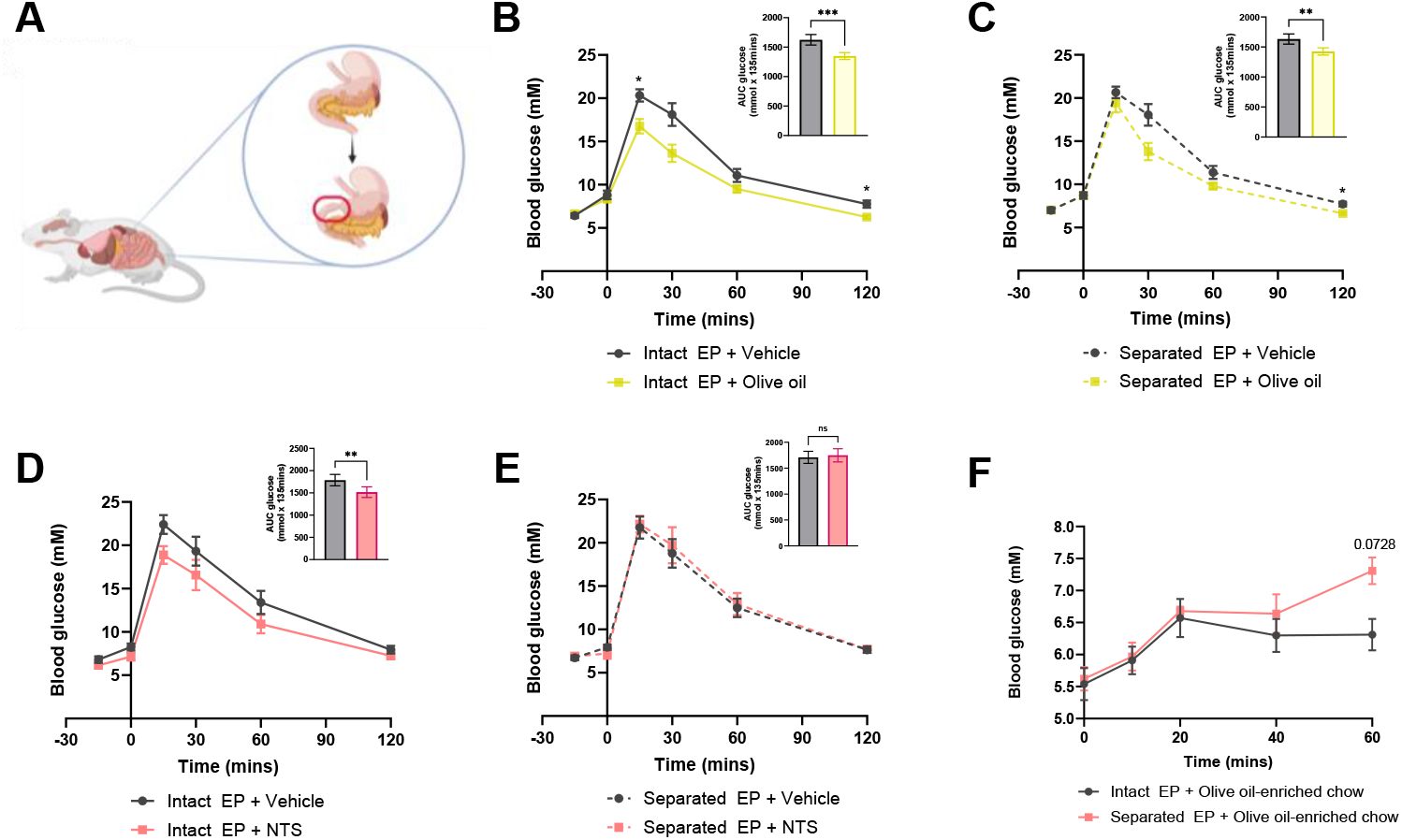
The enteropancreatic axis is necessary for the glucoregulatory effects of neurotensin. Intraperitoneal glucose tolerance tests (IPGTT) were performed in mice that had either undergone surgical separation of the connection between the proximal duodenum and pancreas, or sham surgery. **A).** Schematic depicting surgical separation of proximal duodenum and pancreas in mice. **B).** IPGTT with olive oil or vehicle oral gavaged (OG)-15 minutes prior to glucose load in mice with an intact connection between the proximal duodenum and pancreas. n=15/group. Inset: AUC of blood glucose over IPGTT **C).** IPGTT with olive oil or vehicle OG-15 minutes prior to glucose load in mice with a severed connection between the proximal duodenum and pancreas. n=15/group. Inset: AUC of blood glucose over IPGTT **D).** IPGTT with intraperitoneal (IP) neurotensin or vehicle 15 minutes prior to glucose load in mice with an intact connection between the proximal duodenum and pancreas. n=16/group. Inset: AUC of blood glucose over IPGTT. **E).** IPGTT with IP neurotensin or vehicle 15 minutes prior to glucose load in in mice with severed connection between the proximal duodenum and pancreas. n=16/group. Inset: AUC of blood glucose over IPGTT. **F).** Fast-refeed study with mice with an intact or severed connection between the proximal duodenum and pancreas. Mice were refed olive-oil enriched chow at t0 and blood glucose monitored for 1h. n=8/group. All IPGTT data were analysed with 2-way ANOVA with Tukey’s post-hoc analysis test. AUC data were analysed with paired t-test.

**Figure 4.**
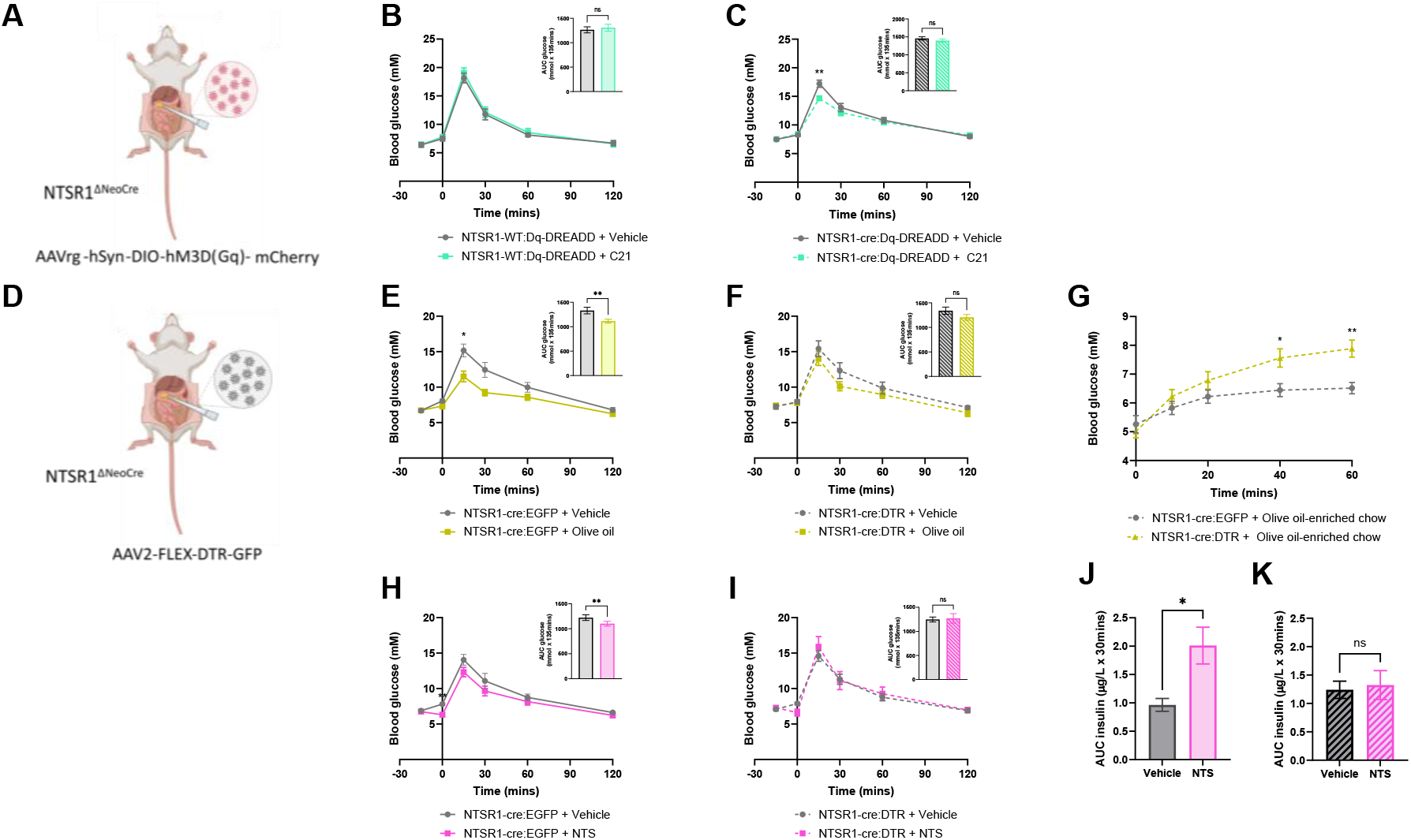
Enteropancreatic-NTSR1 neurons are sufficient and necessary for the effects of olive oil and neurotensin on glucose tolerance. *NTSR1-WT* and *NTSR1-cre* mice were injected with AAVrg-hSyn-DIO-hM3D(q) or of AAV2-FLEX-DTR-GFP into the pancreas and intraperitoneal glucose tolerance tests (IPGTT) performed two weeks later or one week after injection of diphtheria toxin, respectively. **A).** Schematic depicting injection of AAVrg-hSyn-DIO-hM3D(q) into the pancreas. **B).** IPGTT with intraperitoneal (IP) vehicle or C21 IP 15 minutes prior to glucose load in *NTSR1-WT* mice. n=4/group. Inset: AUC of blood glucose over IPGTT. **C).** IPGTT with IP C21 or vehicle 15 minutes prior to glucose load in *NTSR1-cre* mice. n=17/group. Inset: AUC of blood glucose over IPGTT **D).** Schematic depicting injection of AAV2-FLEX-DTR-GFP into the pancreas. **E).** IPGTT with oral gavaged (OG) olive oil or vehicle 15 minutes prior to glucose load in *NTSR1-cre* mice injected with AAV2-pCAG-FLEX-EGFP-WPRE. n=10/group. Inset: AUC of blood glucose over IPGTT **F).** IPGTT with OG olive oil or vehicle 15 minutes prior to glucose load in *NTSR1-cre* mice injected with AAV2-FLEX-DTR-GF. n=10/group. Inset: AUC of blood glucose over IPGTT **G).** Fast-refeed study with *NTSR1-cre* mice injected with either AAV2-pCAG-FLEX-EGFP-WPRE or AAV2-FLEX-DTR-GFP. Mice were refed olive-oil enriched chow at t0 and blood glucose monitored for 1h. n =11-12/group. **H).** IPGTT with IP neurotensin or vehicle 15 minutes before glucose load in *NTSR1-cre* mice injected with AAV2-pCAG-FLEX-EGFP-WPRE. n=10/group. Inset: AUC of blood glucose over IPGTT **I).** IPGTT with IP neurotensin or vehicle 15 minutes before glucose load in *NTSR1-cre* mice injected with AAV2-FLEX-DTR-GFP. n=10/group. Inset: AUC of blood glucose over IPGTT **J).** Plasma insulin levels measured 30 minutes post-IP neurotensin or vehicle administration and 15 minutes post-intraperitoneal glucose load (t30 on IPGTT) in *NTSR1-cre* mice injected with AAV2-pCAG-FLEX-EGFP-WPRE. n=6/group. **K).** Plasma insulin levels measured 30 minutes post-IP neurotensin or vehicle administration and 15 minutes post-intraperitoneal glucose load (t15 on IPGTT) in *NTSR1-cre* mice injected with AAV2-FLEX-DTR-GFP. n=6/group. All IPGTT and fast-refeed data was analysed with 2-way ANOVA with Sidak’s post-hoc analysis test and embedded AUC data were analysed with paired t-test. Plasma insulin data was analysed with a paired t-test.

Having identified that the NTSR1 is expressed in neurons of the myenteric plexus of the proximal duodenum, we assessed whether neurotensin could activate these neurons in a NTSR1-dependent manner. We isolated myenteric neurons (MENs) from the proximal duodenum of *Ntsr1-cre:tdTomato* mice. We found cultured MENs expressing tdTomato (NTSR1+) to be activated by neurotensin; this effect was blocked by an NTSR1 antagonist (Fig.2C&D). Using *Ntsr1-cre:tdTomato* reporter mice we found that NTSR1-expressing neurons extended from the myenteric plexus in the duodenum and extended in fibre bundles into the pancreas (Fig. 2E), where individual NTSR1-expressing processes could be visualised close to pancreatic islets (Fig. 2F). Following injection of a retrograde AAV encoding enhanced green fluorescent protein (eGFP) into the pancreas, we were able to detect eGFP expression in the myenteric plexus of the duodenum (Fig. 2G).

### An intact enteropancreatic connection is required for the glucoregulatory effects of neurotensin and olive oil

To determine the necessity of these enteropancreatic neurons for the effects of olive oil and neurotensin on glucose tolerance, the proximal duodenum was partially surgically separated from its adjacent pancreatic section (Fig.3A). An eGFP-encoding AAV (AAV5.hSyn.eGFP.WPRE.bGH) was injected into the pancreas adjacent to the proximal duodenum to allow verification that enteropancreatic neuronal innervation of the pancreas was reduced; a significant reduction in eGFP expression was observed in the duodenum of mice with a separated enteropancreatic connection (Extended Data Fig.5B). Following recovery, mice underwent a series of glucose tolerance tests. All mice exhibited significantly improved glucose tolerance when treated with the GLP-1R agonist exendin-4 or with an oral bolus of whey protein, suggesting the pancreas remained functional and responsive to a non-lipid nutrient stimulus and a systemic gut hormonal signal (Extended Data Fig.5C-F). In contrast, both neurotensin and olive oil significantly improved glucose tolerance in mice with an intact enteropancreatic connection (Fig.3B-E) but not in mice in which the enteropancreatic neurons were disrupted. When fasted mice were refed olive–oil enriched chow, mice with a disrupted enteropancreatic connection had a significantly higher postprandial glucose excursion compared to control mice (Fig.3F). Together, these data suggest that the glucose lowering effects of olive oil ingestion and neurotensin administration are reliant on enteropancreatic neurons originating in the proximal duodenum.

### NTSR1-expressing enteropancreatic neurons are sufficient to improve glucose tolerance and necessary for the glucoregulatory effect of neurotensin and olive oil

We next investigated whether activating NTSR1-expressing enteropancreatic neurons could modulate glucose tolerance. A cre-dependent retrograde stimulatory DREADD-encoding AAV, (AAVrg-hSyn-DIO-hM3D(Gq)-mCherry) was injected into the head of the pancreas in *NTSR1-cre* mice (Fig.4A). Post-experimental dissection of the longitudinal muscle of the myenteric plexus (LMMP) confirmed mCherry expression in the neurons of the myenteric plexus. Two weeks later, to allow for recovery and sufficient gene expression, intraperitoneal administration of the DREADD ligand compound 21 (C21) 15 minutes prior to an intraperitoneal glucose tolerance test (IPGTT) significantly improved glucose tolerance (Fig.4B&C). Activation of these neurons caused a trend for increased circulating insulin levels (Extended Data Fig.6A-C). In a separate experiment in these mice, administration of C21 did not alter core body temperature, suggesting it was not engaging major central NTSR1 circuits (Extended Data Fig. 6D). These data suggest that activation of NTSR1-expressing neurons that extend to the pancreas from the myenteric plexus of the proximal duodenum is sufficient to improve glucose tolerance.

To assess whether NTSR1-expressing enteropancreatic neurons are necessary for the glucoregulatory effects of peripheral neurotensin and olive oil, a cre-dependent Diphtheria Toxin Receptor-encoding AAV (AAV2-FLEX-DTR-GFP) was injected into the head of the pancreas of NTSR1-cre mice (Figure 4D). As a viral control, the same cre-dependent DTR-encoding AAV (AAV2-FLEX-DTR-GFP) was injected into the pancreas of NTSR1-WT mice and as a genotype control, a cre-dependent GFP-encoding AAV (AAV2-pCAG-FLEX-EGFP-WPRE) was injected into the pancreas of NTSR1-cre mice. Following a 2-week rest period to allow sufficient viral expression, all mice were injected with diphtheria toxin intraperitoneally, which specifically killed NTSR1-expressing enteropancreatic neurons in NTSR1-cre mice injected with AAV-DTR (Extended Data Fig.7B). All mice exhibited significantly improved glucose tolerance when treated with the GLP-1R agonist exendin-4 or with an oral bolus of whey protein, suggesting the pancreas remained functional and responsive to a non-lipid nutrient stimulus and a systemic gut hormonal signal (Extended Data Fig.7C-H). In glucose tolerance tests, both peripheral neurotensin and olive oil retained their glucoregulatory effects in viral and genotype controls (Fig.4E&H, Extended Data Fig.7I&J). However, in NTSR1-cre mice injected with AAV2-FLEX-DTR-GFP, in which enteropancreatic NTSR1-expressing cells had been ablated, neither neurotensin nor olive oil improved glucose tolerance (Fig4.E-F,H-I). In accord with this lack of effect on glucose levels, peripheral neurotensin administration significantly increased circulating insulin levels in genotype control mice but not in NTSR1-cre mice injected with AAV2-FLEX-DTR-GFP (Fig4J&K). NTSR1-cre mice injected with AAV2-FLEX-DTR-GFP also experienced an enhanced glucose excursion compared to genotype control mice in a fast-refeed study where mice were refed olive oil-enriched chow (Fig.4G). These data suggest that NTSR1-expressing enteropancreatic neurons have a role in glucose regulation, stimulating insulin secretion in response to both exogenous neurotensin and physiological neurotensin release following ingestion of olive oil.

## Discussion

Dietary olive oil has previously been linked to improvements in glycaemia and a reduced risk of developing type 2 diabetes in mice and humans^5,42,43^. However, the underlying mechanism for the glucoregulatory effects of olive oil is poorly characterised. Here, we have shown for the first time that ingested olive oil drives an acute improvement in glucose tolerance in mice through driving neurotensin release. We have further demonstrated that this acute glucoregulatory effect of olive oil depends on the digestion of olive oil into its component fatty acids, and is at least partially mediated by the FFAR4.

There is much current interest in agonists of the incretin gut hormones receptors: GLP-1R and GIPR as pharmacotherapies for enhancing glucoregulation. Administration of oleic acid, the major fatty acid constituent of olive oil, has been shown to stimulate release of the gut hormone neurotensin in both rodents and humans^7,10^. Neurotensin has previously been suggested to have a role in glucose regulation, though reported results vary and the mechanisms are unclear^21,22,24^. Previous studies have found neurotensin to have both inhibitory and stimulatory effects on insulin release in vitro and in vivo^21–23,25,44^, though more recent work has suggested a synthetic NTSR1 agonist and a PEGylated form of neurotensin can improve glucose tolerance^26,27^. Here, we provide evidence that neurotensin acts as an incretin and has the potential to improve glucoregulation in both lean and diet-induced obese mice.

Our data shows that neurotensin works to promote insulin secretion via NTSR1-expressing enteropancreatic neurons that originate in the proximal duodenum. Although enteropancreatic neurons were first discovered nearly fifty years ago^45^ they have been relatively little studied. Serotonergic and NADPH diaphorase-expressing enteropancreatic neurons have been reported to have an inhibitory and excitatory role in exocrine pancreas secretions, respectively, and myenteric cholinergic neurons have been shown to provide excitatory inputs to the pancreas^39,40,46^. Using surgical and genetic mouse models, we have demonstrated that activation of NTSR1-expressing enteropancreatic neurons is sufficient to improve glucose tolerance, and these neurons are necessary for the glucoregulatory effects of neurotensin and olive oil. This represents the first functional role for a population of enteropancreatic neurons in the modulation of the endocrine pancreas.

Ingestion and infusion of olive oil and oleate has previously been reported to stimulate endogenous secretion of neurotensin^7,8,10^, and we also found that oral gavage of olive oil increased circulating levels of neurotensin. However, neurotensin has a short half-life in circulation^47^, and it is possible that other agonists of the NTSR1, such as xenin, also play a role in the glucoregulatory effect of olive oil. Xenin is a molecule secreted from enteroendocrine K cells, and exogenous xenin has been reported to potentiate the insulinotropic actions of GIP and bind to the NTSR1 to promote gastric emptying^48^. There are, however, no reports of a physiological role for endogenous xenin.

The NTSR1 has also been found to promote lipid absorption, and knockout of the NTSR1 has been shown to protect rodents from diet-induced obesity^19^. Whilst we have demonstrated that the glucoregulatory effects of exogenous neurotensin and olive oil occur before any increase in circulating triglycerides is detected, chronically agonising the NTSR1 as a therapy for impaired glucoregulation may result in weight gain and obesity. Further research into the enteropancreatic NTSR1 neurons might identify other drug targets to facilitate the more specific targeting of these neurons for glucoregulation. We have also shown that neurotensin can have an additive effect on glucoregulation when administered in combination with the GLP-1 receptor agonist exendin-4, further underpinning the potential of NTSR1-specific therapeutics for the treatment of type 2 diabetes. Chronic studies in disease models are now necessary to determine the therapeutic utility of the NTSR1 system.

This is the first characterisation of a functional role for a neuronal population connecting the gut directly to the pancreas. The position of these neurons in the upper duodenum allows them to respond rapidly to dietary nutrients to drive appropriate pancreatic responses. There may also be additional physiological roles for this enteropancreatic system. Further work is necessary to explore whether NTSR1 agonists and other agents targeting enteropancreatic neurons may be useful in the treatment of metabolic disease.

## Contact for reagent and resource sharing

Further information and requests for reagents may be directed to and will be fulfilled by the Lead Contact, Kevin G Murphy. The data that support the findings of this study are available from the corresponding author upon reasonable request.

## Experimental model and subject details

### Mice

Six- to twelve-week-old male and female C57bl/6 mice were used in studies, unless otherwise stated. *Ntsr1^ΔNeoCre^* (stock no. 033365) were bought in from Jax and breeding pairs set up as hemizygotes x littermate control. *Ntsr1^ΔNeoCre^* mice hemizygous for cre recombinase are noted as *NTSR1-cre*. *Ntsr1^ΔNeoCre^* mice without cre recombinase expression are notated as *NTSR1-WT. Ntsr1-cre:tdTomato* mice were generated by crossing an *Ai9* female (stock no. 007909) with a *NTRS1-cre* male. The *NTSR1^ΔNeoCre^*allele was detected by PCR genotyping with primers 5’-GCTAGAGGGAAACCGTTGTG-3’, 5’-GTTCTCCAGGAAGCCAAACA-3’ and 5’-GAGAGGGATAAGGGGCAAGA-3’ where WT and mutant alleles produced PCR products of 288 bp and 202 bp, respectively. The *Ai9* allele was detected by PCR genotyping with primers 5’-AAGGGAGCTGCAGTGGAGTA-3’, 5’-CCGAAAATCTCTGGGAAGTC-3’, 5’-GGCATTAAAGCAGCGTATCC-3’ and 5’-CTGTTCCTGTACGGCATGG-3’ where WT and mutant alleles produced PCR products of 297 bp and 196 bp, respectively.

GLP-1R fl/fl mice were kindly gifted by Professor Randy Seeley at University of Michigan and crossed with Pdx1-Cre/Esr1 mice (JAX strain: #024968). Tamoxifen (100mg/kg) was injected intraperitoneally for 4-days to induce beta cell knockout of the GLP-1R.

### Primary Cell Culture

Murine myenteric enteric neurons (MENs) were isolated from the proximal duodenum of mice for use in activation studies. 6cm of proximal duodenum were dissected and longitudinal muscle/myenteric plexus (LMMP) carefully peeled off and placed in ice-cold digestion liquid (0.166mg/ml liberase in filtered Krebs-Ringer Bicarbonate Buffer (HKRB).

LMMP were next digested for 45 minutes at 37°^C^, washed in PBS and placed in complete enteric neuronal media (Neurobasal A Media with B-27 supplement, 2mM L-glutamine, 10% FBS, 1% penicillin/streptomycin and 10ng/ml GDNF). MENs were dispersed by trituration using a p200 filter tip and pipette and filtered through a 70µM filter. Released myenteric neurons were then resuspended in complete enteric neuronal media and maintained at 37°^C^, 5% CO_2_ on laminin- and poly-D-lysine-coated slides.

Murine islets were isolated following injection of Collagenase NB 8 broad range (1mg/ml; Universal Biologicals Ltd) into the bile duct of mice. Inflated pancreata were then digested at 37°^C^ following which pancreata underwent a series of washes and manual digestions using cold RPMI (RPMI 1640 medium; Merck, USA) and a Rotina 380R, Hettich Centrifuge. A density gradient was next created using histopaque 1119 (Merck, USA), histopaque 1083 (Merck, USA) and RPMI medium no additives. Isolated islets were then picked using a binocular microscope and maintained at 37°^C^, 5% CO_2_ in complete media (RPMI media, 10% FBS, 1% penicillin/streptomycin).

## Methods

### Animal Experiments

All mice were kept in individually ventilated cages with *ad libitum* access to water and standard laboratory RM1 chow in a temperate, humidity-controlled environment (21-23°^C^) with constant 12h-light/dark cycle (lights on at 7am; lights off at 7pm), unless otherwise stated. Cages and bedding were changed once every week. To create a diet-induced obese model (DIO), mice were fed a high fat diet (60% fat, 20% protein, 20% carbohydrate; D12492, Research Diets, USA) from 6 weeks of age and studies were performed when they were aged 36 weeks. Mice were monitored daily for their health status. All procedures were carried out according to the Animals (Scientific Procedure) Act, 1986 and Amendment Regulations, 2012 and approved by the Imperial College London Animal Welfare and Ethical Review Body.

### Nodose Ganglion Surgery

6-week-old C5BL6/J mice were purchased from Charles River, U.K. and left to acclimatise for 1 week. Following this, mice were injected with either NTS-saporin [IT-56] or BLANK-saporin [IT-21] (1.5µg/µl; 0.5µl / nodose ganglia; Advanced Targeting Systems, CA) into the nodose ganglia. Injections were performed unilaterally with one week’s rest before the second ganglion was injected. All injections were performed under isoflurane anaesthesia and mice were administered buprenorphine (0.12mg/kg) and carprofen (4mg/kg) as the perioperative analgesia. Mice were also administered atropine sulphate (2µg/kg) perioperatively to help prevent cardiac arrest caused by vagal manipulation during surgery. Postoperatively, mice were maintained on a liquid diet for 3 days (Ensure Plus Vanilla, 1.5kCal/ml) to minimise risk of unexpected gastrointestinal disturbances following vagal manipulation and carprofen was added to their drinking water (2.72mg/100ml).

### Intracerebroventricular (ICV) Cannulation

6-week-old C57bL6/J mice were purchased from Charles River, U.K. and left to acclimatise for 1 week. Following this, mice were implanted with a stainless-steel cannula into the 3^rd^ ventricle (−0.8mm anterior/posterior, +0.1mm lateral, −2.5mm dorsal/ventral). All implantations were performed under isoflurane anaesthesia and mice were administered buprenorphine (0.12mg/kg) and carprofen (4mg/kg) as the perioperative analgesia. Postoperatively, carprofen was added to drinking water (2.72mg/100ml) for 3 days.

All injections (0.5µl/injection) were performed at a maximum infusion rate of 30µl/hour using a locking internal cannula with 0.5mm projection in conscious animals. Cannula were left indwelling for an additional minute post-injection to prevent backflow with withdrawal.

### *In Vivo* AAV Injections

8-week-old *NTRS1-cre* and *NTSR1-WT* mice were injected with AAVrg-hSyn-DIO-hM3D(Gq)-mCherry (7×10^12^ vg/ml; 1µl/injection), AAV2-FLEX-DTR-GFP (5×10^12^ vg/ml; 1µl/injection), AAV2-pCAG-FLEX-EGFP-WPRE (7×10^12^ vg/ml; 1µl/injection), AAV5.hSyn.eGFP.WPRE.bGH (2.2×10^13^ vg/ml; 1µl/injection), pAAVrg-hSyn-EGFP (7×10¹² vg/mL; Addgene #50465-AAVrg; 1 μL/injection) or pAAV9-hSyn-EGFP (7×10¹² vg/mL; Addgene #50465-AAV9; 1 μL/injection) in three separate areas in the head of the pancreas. All injections were performed under isoflurane anaesthesia and mice were administered buprenorphine (0.12mg/kg) and carprofen (4mg/kg) as the perioperative analgesia. Postoperatively, carprofen was added to drinking water (2.72mg/100ml) for 3 days and jelly infused with buprenorphine (0.1mg/ml) was placed in cages twice daily.

### Diphtheria Toxin-Mediated Ablation of NTSR1-Expressing Cells

*NTRS1-cre* mice injected with AAV2-FLEX-DTR-GFP or AAV2-pCAG-FLEX-EGFP-WPRE and *NTSR1-WT* injected with AAV2-pCAG-FLEX-EGFP-WPRE received diphtheria toxin (30ng/g, Sigma, UK) via intraperitoneal injection once daily for two days. A week later, mice underwent intraperitoneal glucose tolerance tests.

### Surgical Separation of the Enteropancreatic Connection

7-week-old C57bL6/J mice were purchased from Charles River and left to acclimatise for 1 week. To achieve surgical separation of the enteropancreatic network, the connection between the first 6 cm of the proximal duodenum and adjacent pancreas was severed through blunt dissection using forceps, or left intact in control mice. Prior to separation, the pancreatic duct was located and care was taken to preserve it during surgery and all mice received pancreatic injection of AAV5.hSyn.eGFP.WPRE.bGH for model validation. All surgeries were performed under isoflurane anaesthesia and mice were administered buprenorphine (0.12mg/kg) and carprofen (4mg/kg) as perioperative analgesia. Postoperatively, carprofen was added to drinking water (2.72mg/100ml) for 3 days.

### Measurement of Olive Oil induced Neurotensin Secretion

Mice were fasted for 5h with ad libitum access to water before being administered olive oil (200µl/mouse; Napolina) or control (dH20) by oral gavage. Mice were sacrificed 5 minutes after gavage and blood samples collected in EDTA-coated microvettes and spun at 3,600xg for 15 minutes. Plasma samples were then stored at −80°^C^ until analysis by ELISA.

### Intraperitoneal Glucose Tolerance Tests

Tests were performed on mice following a 5h fast with *ad libitum* access to water. Following a baseline blood glucose measurement, mice were challenged with glucose (20%, 2g/kg) IP and changes in blood glucose measured over the following 120 minutes. Blood glucose readings were taken from a 0.5cm tail-tip venesection and measured using an AccuCheck glucometer and glucose strips. Blood samples for plasma insulin and triglyceride analysis were collected in EDTA-coated microvettes and spun at 3,600xg for 15 minutes. Plasma samples were then stored at −80°^C^ until analysed.

For assessment of the role of different types of oil on glucose tolerance, olive oil (200µl/mouse; Napolina) and coconut oil (200µl/mouse; VITA COCA) or control (dH20) were administered via oral gavage 15 minutes before glucose IP. The NTSR1 antagonist SR48692 (80µg/kg, Tocris) or vehicle (1% DMSO) was administered intraperitoneally 60 minutes prior to glucose IP. Whey protein (10% w/v; 300µl/mouse; Pure Whey Isolate 90, Bulk Powders, UK) or vehicle (dH20) was administered via oral gavage 15 minutes before glucose IP. Neurotensin acetate salt (6mg/kg, Bachem) or vehicle (dH20) was administered intraperitoneally 15 minutes prior to glucose IP. Exendin-4 (0.6µg/kg, Wuxi Apptec) was co-administered intraperitoneally with glucose. 4DAMP (1µg/kg) or vehicle (1% DMSO) was administered intraperitoneally 15 minutes prior to glucose IP. Compound 21 (0.3mg/kg, HelloBio) or vehicle (saline) was administered intraperitoneally 15 minutes prior to glucose IP. Orlistat (250mg/g of fat, Cambridge Biosciences), dissolved in olive oil, or vehicles (olive oil, dH20) were administered via oral gavage 15 minutes prior to glucose IP and AH-7614 (5mg/kg, Tocris) or vehicle (10% DMSO in corn oil) administered subcutaneously 15 minutes prior to olive oil oral gavage.

### Fast-Refeeding Study

Tests were performed on mice following a 16h fast with *ad libitum* access to water. Following the fast, olive-oil enriched RM1 chow (2ml olive oil: 1 (∼3g) pellet RM1 chow; 1 pellet / animal in cage) was given to the mice. Blood glucose readings were taken from a 0.5cm tail-tip venesection and measured using an AccuCheck glucometer and glucose strips and change in blood glucose measured over the following 60 minutes. To investigate the role of NTSR1, SR48692 (80µg/kg, Tocris) or vehicle (1% DMSO) were administered intraperitoneally 60 minutes prior to the return of olive-oil enriched chow.

### Food Intake Study

Tests were performed on individually caged mice following a 12h fast with *ad libitum* access to water in the light phase. SR48692 (2.5µg; Tocris) or vehicle (1% DMSO; Sigma-Aldrich) was administered ICV 20 minutes prior to intraperitoneal injection of neurotensin (6mg/kg, Bachem) or vehicle (dH20). RM1 chow was weighed and returned to cages and food intake measured over the following 1h.

### Core Temperature Study

Tests were performed on mice following a 5h fast with *ad libitum* access to water. Rectal temperature was taken using a RS PRO RS41 K handheld digital thermometer with external K-type thermocouple bead probe and measured at baseline and 15 minutes post-treatment.

Neurotensin (6mg/kg, Bachem) or vehicle (dH20) were administered ICV into the 3^rd^ ventricle. Compound 21 (0.3mg/kg, HelloBio) or vehicle (saline) were administered intraperitoneally.

### Metabolic Assays

#### Plasma Insulin Assay

Plasma insulin concentration from collected plasma samples were assayed using the Mouse Insulin ELISA (Mercodia, Sweden) according to the manufacturer’s instructions.

#### Plasma Triglyceride Assay

Plasma triglyceride concentrations in collected plasma samples were assayed using the Triglycerides TR210 Assay Kit (Randox, UK) according to the manufacturer’s instructions.

#### Plasma Neurotensin Assay

Plasma neurotensin concentration from collected plasma samples were assayed using the Mouse Neurotensin ELISA (E229Mo, BT Labs) according to the manufacturer’s instructions.

### RNA Isolation and Gene Expression Analysis by qPCR

#### Preparation of Tissue Samples for RNA Extraction

For the quantification of NTSR1 gene expression in different glucoregulatory tissues, punch biopsies of arcuate nucleus were made using the optic chiasm and third ventricle as landmarks and nodose ganglia were carefully dissected from mice and snap frozen at −80°^C^. Isolated islets and myenteric enteric neurons were isolated as described above and snap frozen at −80°^C^.

#### RNA isolation

Total RNA was isolated from tissues and cells using TRIsure (Bioline) and homogenised using a TissueLyser II (Qiagen, Hildon). Samples underwent a series of separation steps involving chloroform and centrifugation, following which RNA was precipitated and visualised using GlycoBlue (Invitrogen). The amount and purity of isolated RNA was assessed using a Nanodrop Lite Spectrophotometer (Thermofisher Scientific, Madison). Genomic DNA was removed from samples using RNase-free DNase (Promega) following which 1µg of RNA was reverse-transcribed using a High Capacity cDNA Reverse Transcription Kit (Applied Biosystems), according to the manufacturer’s instructions.

#### Quantitative Real-Time PCR (qPCR)

qPCR was performed using Maxima Probe Master Mix (Life Technologies, Paisley) in a C1000 CFX384 Real-Time System Thermocycler using Taqman PCR core reagents (Applied Biosystems). To quantitatively analyse NTSR1 expression, the taqman probe for NTSR1 was used (assay ID: Mm00444459_m1; Applied Biosystems) and results analysed using the comparative C_T_ method. Relative NTSR1 expression was normalised to the ‘housekeeping’ gene β-actin mRNA (assay ID: Mm02619580_g1; ThermoFisher).

#### Tissue Clearing and Imaging

Mice were given an overdose of pentobarbital and transcardially perfused with 0.01M PBS followed by 4% PFA. The first 6 cm of the proximal duodenum and its adjacent pancreatic section were dissected out and fixed overnight in 4% PFA at 4°C. Tissues were cleared using RTF tissue clearing^1^ to retain endogenous fluorescence. Briefly, tissue was sequentially immersed in 3 different solutions constituted by triethanolamine (TEA) (Sigma) and formamide (F) (ThermoFisher) and dH20 (W); RTF-R1 (30%TEA/40%F/30%W), RTF-R2 (60%TEA/25%F/15%W), RTF-R3 (70%TEA/15%F/15%W) over 1 day. The cleared sample was then imaged on an inverted confocal Leica Stellaris 8 microscope (Facility for Imaging by Light Microscopy, Imperial College London) and further processed in ImageJ. iDISCO whole-mount immunolabelling^2^ without methanol pre-treatment was conducted prior to tissue clearing and imaging. To label insulin-secreting beta cells within the pancreas, a guinea pig anti-insulin antibody was used (1:100; ab7842, Abcam) and immunofluorescence was identified using a goat anti-guinea pig secondary IgG antibody conjugated to Alexa-Fluor 647 (1:500; ThermoFisher).

#### LMMP Immunohistochemistry

NTSR1-cre:tdTomato mice injected with AAV2-FLEX-DTR-GFP, AAV5.hSyn.eGFP.WPRE.bGH or AAV2-pCAG-FLEX-EGFP-WPRE-injected mice were killed by cervical dislocation and the LMMP of the proximal duodenum carefully dissected out. LMMP was fixed overnight in 4% paraformaldehyde (PFA, Sigma-Aldrich, UK). To identify NTSR1-expression in the myenteric plexus, LMMP were stained with the pan-neuronal marker polyclonal rabbit anti-beta tubulin III (1:200; ab18207; Abcam, UK), mouse anti-beta tubulin III (1:1000; ab78078; Abcam) or chicken anti-GFP (ab13970, Abcam, UK; 1:1000). All samples were blocked in 0.1M PBS with 10% neonatal goat serum (New Zealand Origin; Gibco), 4% bovine albumin serum (BSA, Sigma-Aldrich) and 0.3% triton-X (Sigma-Aldrich) for 1 hour at room temperature prior to primary antibody, and then overnight at 4°^C^. Immunofluorescence was identified using a goat anti-rabbit secondary IgG antibody conjugated to AlexaFluor-488 (1:500; ThermoFisher), goat anti-mouse secondary IgG antibody conjugated to AlexaFluor-488 (1:500; ThermoFisher) or goat anti-rabbit secondary IgG antibody conjugated to AlexaFluor-633 (1:500; ThermoFisher). Nuclei were stained using DAPI mounting media (ProLong™ Diamond Antifade Mountant with DAPI; ThermoFisher) and images acquired using a Zeiss AxioObserver widefield microscope with x5, x10 or x20 objective lens or confocal microscope Leica Stellaris 8 inverted (Leica) at 40× magnification. Images were analysed in ImageJ.

The neuronal nature of cultured myenteric enteric neurons was confirmed using the technique as described above.

#### Verification of Surgical Separation of the Proximal Duodenum and the Pancreas Model

To validate enteropancreatic separation, the percentage eGFP reflecting virus (intrapancreatically injected AAV5.hSyn.eGFP.WPRE.bGH) expression within total neurons in the LMMP was quantified and compared to control mice. Dissected LMMPs were stained with anti-beta tubulin III and imaged at 20x magnification. In ImageJ, a ‘neuronal’ (beta-tubulin III-expressing) mask and a ‘viral’ (eGFP-expressing) mask was created. The masks were overlaid and the percentage virus within the neuronal mask quantified. Thresholding was kept consistent across channels between mice. Each N value reflects quantification of 2-8 averaged image quantifications for each mouse.

#### Verification of AAV-DTR Ablation Model

To validate ablation of the NTSR1-expressing enteropancreatic cells, the percentage tdTomato, reflecting NTSR1, expressing neurons, within total neurons in the LMMP was quantified and compared between *Ntsr1-cre:tdTomato* mice intrapancreatically injected with AAV2-FLEX-DTR-GFP or AAV2-pCAG-FLEX-EGFP-WPRE. Dissected LMMPs were stained with anti-beta tubulin III and imaged at 10x magnification. In ImageJ, a ‘neuronal’ (beta-tubulin III-expressing) mask and a ‘tdTomato’ (eGFP-expressing) mask was created. The masks were overlaid and the percentage tdTomato within the neuronal mask quantified. Thresholding was kept consistent across channels between mice. Each N value reflects quantification of 2-6 averaged image quantifications for each mouse.

### Calcium activation studies

#### Signalling Assay

To investigate treatment-stimulated cell activation, myenteric enteric neurons were cultured as described above and imaged on day 5 when neuronal projections had begun to form. Changes in intracellular calcium levels were measured using Fluo-4AM Direct labelling (Invitrogen), according to manufacturer’s protocol. Agonist-dependent calcium mobilization was assessed using a Zeiss AxioObserver widefield microscope with a x20 objective lens, capturing 1 frame per second.

Cells were incubated in Fluo-4 AM Direct at 37°^C^ for 30 minutes, followed by a 30-minute incubation at room temperature. Baseline readings were then taken for 60 seconds prior to addition of treatment in a 1:5 dilution (200pM or 100nM neurotensin, Bachem). SR48692 (10µM; Tocris) was added 20 minutes prior to baseline readings. KCl (50mM) was added 5 minutes post-treatment to confirm cell viability.

#### Calcium Imaging Analysis

Images were analysed using ImageJ and a macro package from Imperial FILM facility (Intensity_2). Regions of interest (ROI) were drawn around all obvious cell bodies to capture fluorescence (f). Average fluorescence for baseline readings of each cell were calculated, using which fluorescent change for each cell was quantified (fluorescence at every frame – baseline average / baseline = Δf/f). Max Δf/f over the 2.5-minute period following addition of treatment was calculated and cells stratified into responders and non-responders (responder: max Δf/f of treatment > max Δf/f of baseline + 2x SD; Non-responder: max Δf/f of ligand < max Δf/f of baseline + 2x SD).

#### Quantification and Statistical Analysis

Data were analysed using Prism 9.0 software (GraphPad Software) and expressed as mean ± SEM. Analysis of data with one independent variable was analysed with one-way ANOVA with Tukey’s multiple comparison post hoc test. Data with only two independent variables alongside a continuous outcome variable were analysed using a two-way ANOVA with Sidak’s multiple comparison post hoc test. In studies comparing three or more groups, Tukey’s multiple comparison post hoc test was used. For AUC comparisons between two groups, data were analysed using a paired t-test. Statistical analysis of area under the curve data sets was conducted using a student’s t-test (two-sided). p values <0.05 were considered statistically significant.

## Acknowledgements

The Section of Endocrinology and Investigative Medicine is funded by grants from the MRC, BBSRC, NIHR and is supported by the NIHR Biomedical Research Centre Funding Scheme and the NIHR/Imperial Clinical Research Facility. The views expressed are those of the authors and not necessarily those of the MRC, BBSRC, the NHS, the NIHR or the Department of Health. KGM is supported by Diabetes UK (18/0005886, 20/0006295), the BBSRC (BB/W001497/1, BB/X017273/1), the MRC (MR/Y013980/1) and the Wellcome Trust (310835/Z/24/Z). BYHL and GSHY are supported by BBSRC Project Grant (BB/S017593/1) and the MRC Metabolic Diseases Unit (MC_UU_00014/1). Graphical abstract, Fig. 3A, 4A&D, Extended Data Fig. 2B, 3A, 4A, 5A, 7A were created using BioRender (https://www.biorender.com). We thank Bryn Owen for his comments on the manuscript.

## Author Contributions

A.G.R., L.M., J.L, M.N., P.P., A.M-A., G.D., S.C., C.D., Y.T., T.S-C., A.B.D., A.C., K.G.M. performed experiments, analysed data and/or interpreted data. B.L., A.H., B.J., G.Y., V.S., K.G.M. contributed specific reagents or methods, interpreted data and provided input on the experiments. A.G.R., L.M., K.G.M. wrote the manuscript with input from all co-authors. K.G.M. directed and supervised the study.

---

**Figure S1.**
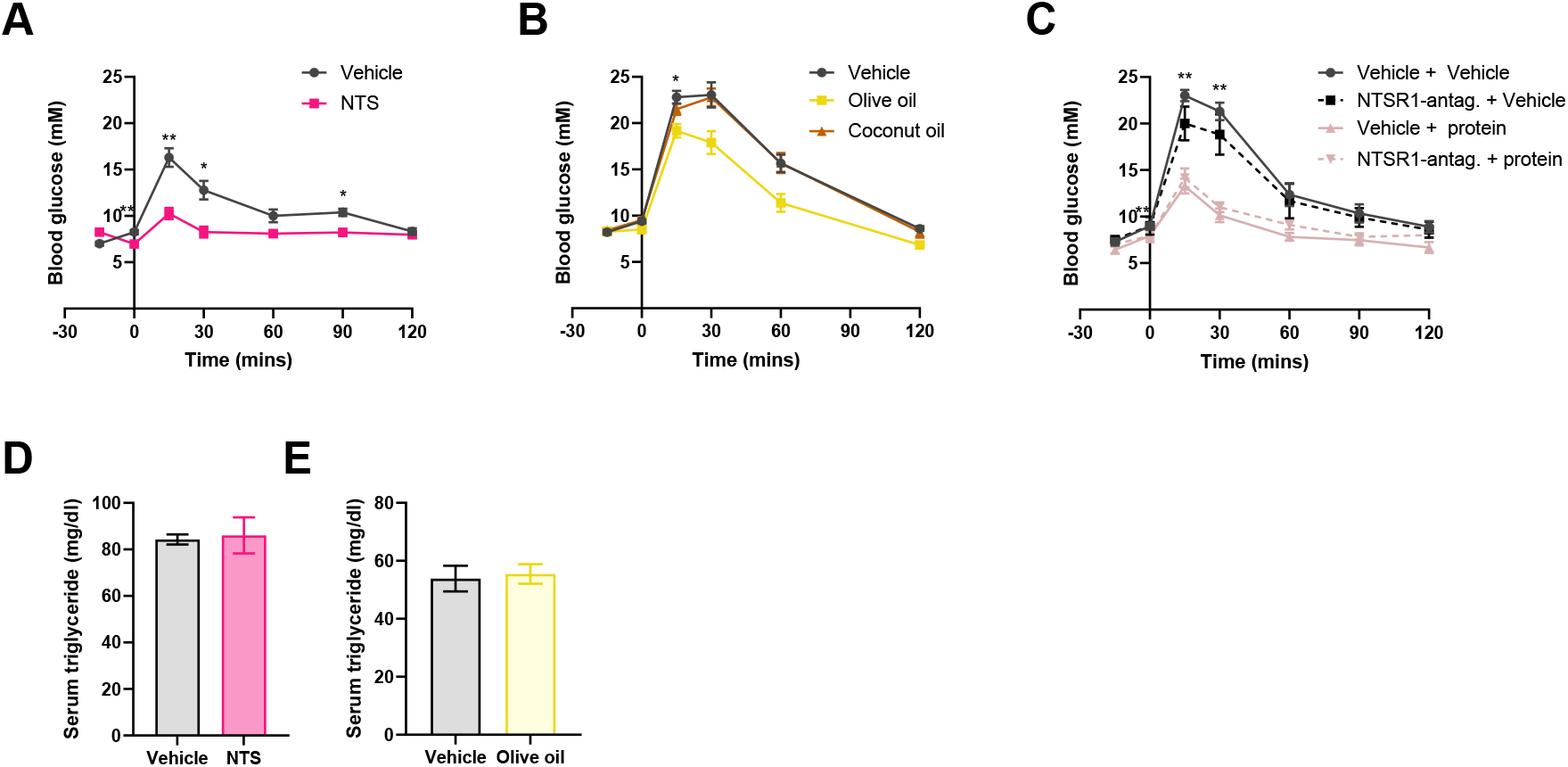
The glucoregulatory effects of olive oil are not replicated by coconut oil and blocking the NTSR1 does not prevent the glucoregulatory effects of another macronutrient. **A).** Oral glucose tolerance test with IP neurotensin or vehicle 15 minutes prior to glucose load. n=8/group. **B).** Intraperitoneal glucose tolerance test (IPGTT) with olive oil, coconut oil or vehicle orally gavaged (OG)-15 minutes prior to glucose load. n=14/group. **C).** IPGTT with 1h IP pretreatment of an NTSR1-specific antagonist followed by OG whey protein (10% weight/vol) or vehicle. n=5/group. **D).** Serum triglyceride levels measured 15 minutes post-IP neurotensin or vehicle administration and intraperitoneal glucose load. n=3/group. **E).** Serum triglyceride levels measured 15 minutes post-OG olive oil or vehicle administration and intraperitoneal glucose load. n=7/group. All IPGTT data was analysed with 2-way ANOVA with Sidak’s post-hoc analysis test. # indicates significance between vehicle and olive oil, $ indicates significance between vehicle + vehicle and vehicle + protein.

**Figure S2.**
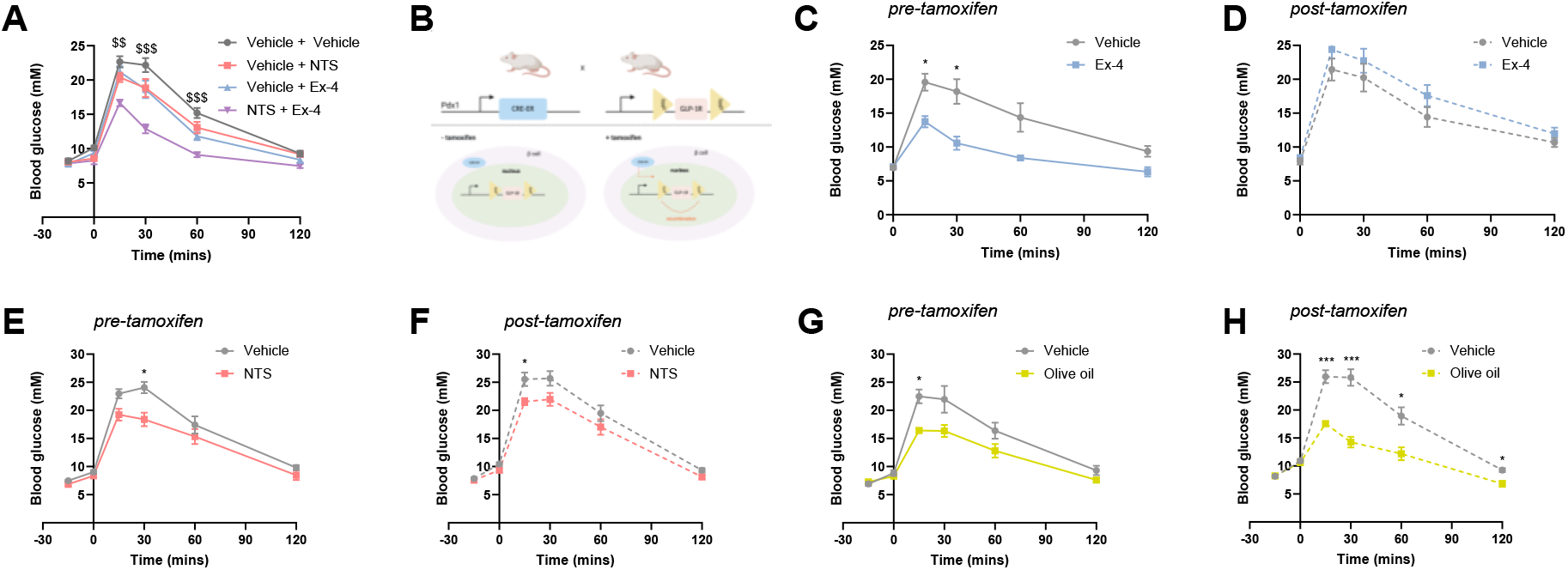
The glucoregulatory effects of neurotensin and olive oil are not mediated by the GLP-1 receptor. **A).** Intraperitoneal glucose tolerance test (IPGTT) with intraperitoneal (IP) neurotensin, exendin-4 (Ex-4) or neurotensin + Ex-4 15 minutes prior to glucose load. n=10/group. **(B).** Schematic of tamoxifen-induced knock out of GLP-1R from pancreatic beta cells using Pdx1CreERT-GLP-1Rfl/fl mice. **C)**. IPGTT in PDX4 mice with olive oil or vehicle 15 minutes prior to glucose load pre-tamoxifen induction. n= 5-7/group. **D).** IPGTT in Pdx1CreERT-GLP-1Rfl/fl mice with olive oil or vehicle 15 minutes prior to glucose load post-tamoxifen induction. n= 5-8/group. **E).** IPGTT in Pdx1CreERT-GLP-1Rfl/fl mice with IP neurotensin or vehicle 15 minutes prior to glucose load pre-tamoxifen induction. n= 9/group. **F).** IPGTT in Pdx1CreERT-GLP-1Rfl/fl mice with IP neurotensin or vehicle 15 minutes prior to glucose load post-tamoxifen induction. n= 16/group. **G).** IPGTT in Pdx1CreERT-GLP-1Rfl/fl mice with Ex-4 or vehicle 15 minutes prior to glucose load pre-tamoxifen induction. n= 6-/group. **H).** IPGTT in Pdx1CreERT-GLP-1Rfl/fl mice with Ex-4 or vehicle 15 minutes prior to glucose load post-tamoxifen induction. n= 6-7/group. All IPGTT data was analysed with 2-way ANOVA with Sidak’s post-hoc analysis test. $ indicates significance between vehicle + vehicle and NTS + Ex-4.

**Figure S3.**
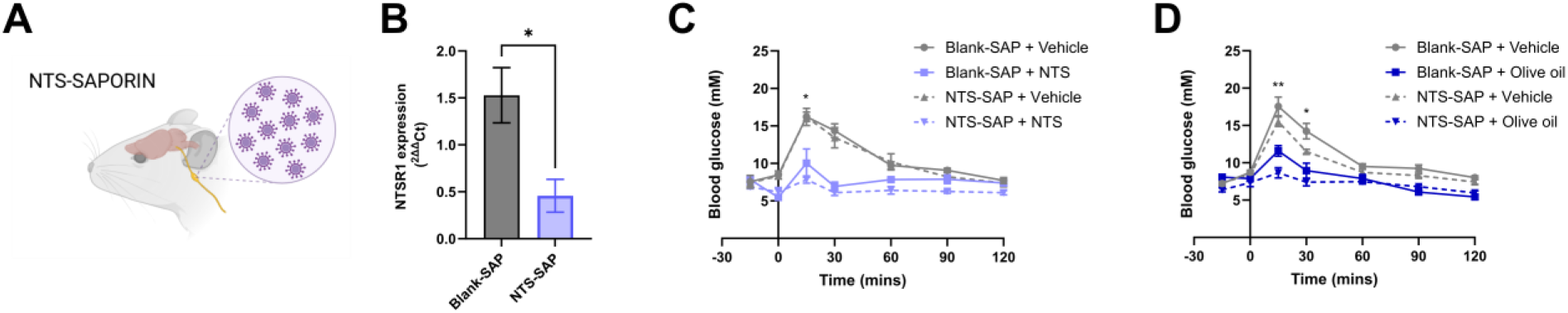
Neurotensin does not improve glucose tolerance via activation of vagal-NTSR1. **A).** Schematic depicting bilateral injection of NTS-saporin or BLANK-saporin into the vagal nodose ganglia of C57BL6/J mice. **B).** qPCR analysis of relative NTSR1 expression in the vagal nodose ganglia of NTS- and BLANK-saporin injected mice. n=4-5/group. **C).** IPGTT with 15-minute pretreatment of IP neurotensin or vehicle 15 minutes prior to glucose load in NTS- and BLANK-saporin injected mice. n=4-6/group. **D).** IPGTT with 15-minute oral gavage pretreatment of olive oil or vehicle 15 minutes prior to glucose load in NTS- and BLANK-saporin injected mice. n=4-6/group. qPCR data analysed using unpaired t-test. IPGTT data analysed using 2-way ANOVA with Tukey’s post-hoc analysis. # indicates significance between NTS-SAP + vehicle and NTS-SAP + NTS or NTS-SAP + vehicle and NTS-SAP + olive oil.

**Figure S4.**
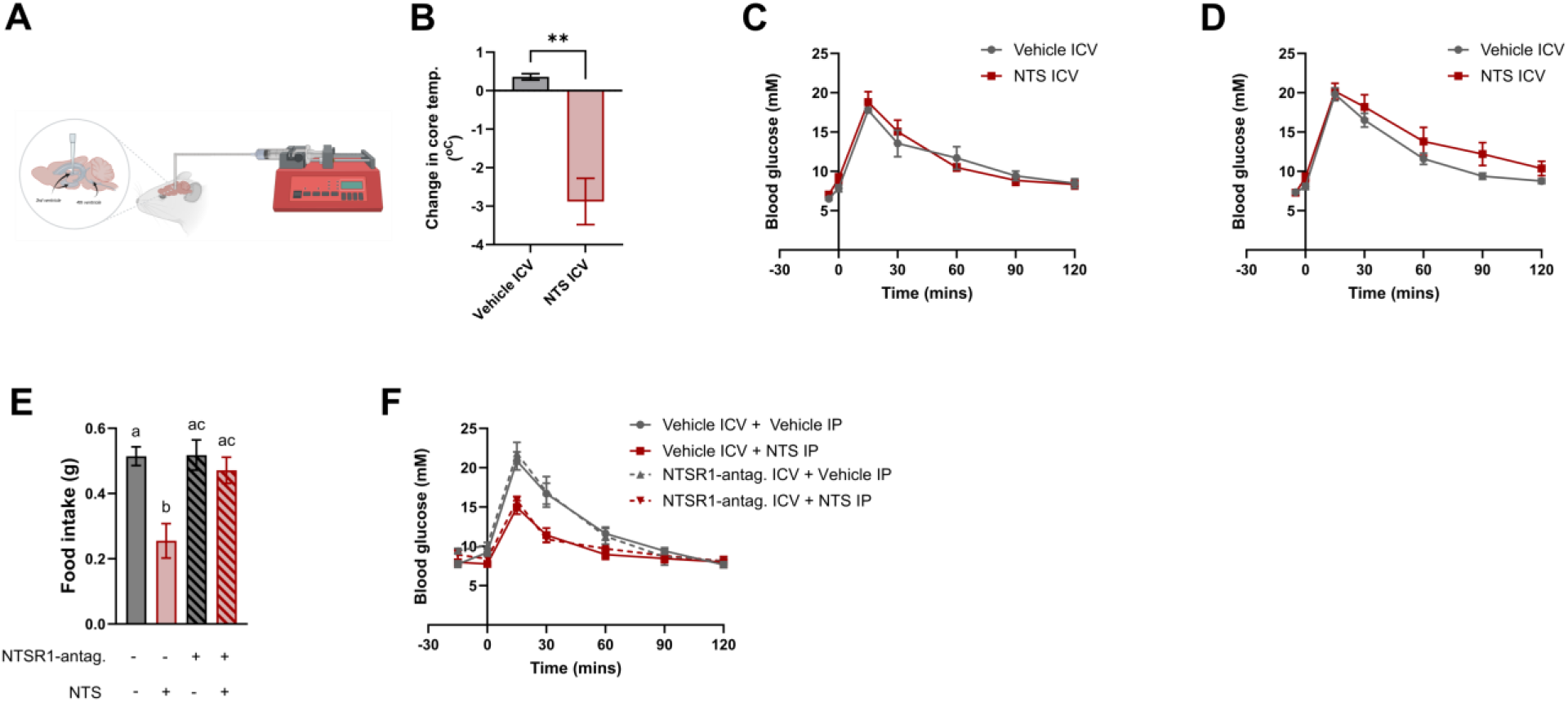
Neurotensin does not improve glucose tolerance via activation of central-NTSR1. **A).** Schematic depicting cannulation into the 3^rd^ ventricle of C57BL6/J mice. **B).** Change in core temperature following central infusion of neurotensin or vehicle into the 3^rd^ ventricle. n=5/group. **C).** Intraperitoneal glucose tolerance test (IPGTT) with low dose (10ng) central administration of neurotensin or vehicle 5 minutes prior to glucose load. n=8-10/group. **D).** IPGTT with high dose (400ng) central neurotensin or vehicle 5 minutes prior to glucose load. n=8-10/group. **E).** Following a 12h fast, mice were centrally injected with an NTSR1 antagonist (SR48692, 2.5µg/mouse) or vehicle followed by peripheral neurotensin (6mg/kg) or vehicle. Food was returned after 10 minutes and food intake was measured 1h later. **F).** IPGTT with 1h pretreatment of central NTSR1-specific antagonist or vehicle followed by intraperitoneal neurotensin or vehicle 15 minutes prior to glucose load. n=4-5/group. Change in core temperature was analysed using an unpaired student’s t-test. Food intake and IPGTT data were analysed using a 2-way ANOVA with Tukey’s post-hoc analysis. # indicates significance between NTSR1-antagonist ICV + vehicle IP and NTSR1-antagonist ICV and NTS IP.

**Figure S5.**
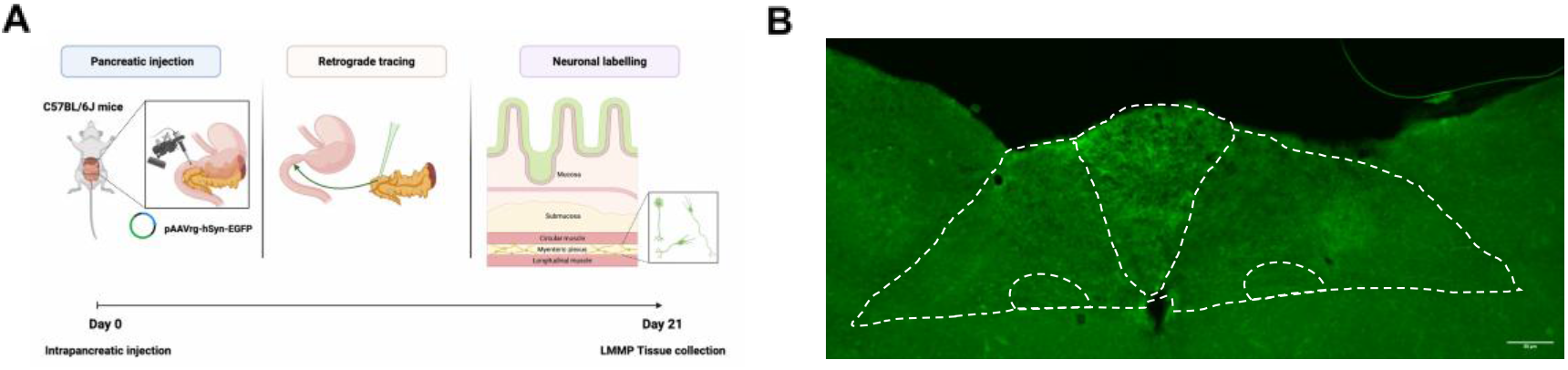
Validation of enteropancreatic-specificity of retrograde tracing. **A).** Schematic depicting microinjection of retrograde tracer into the pancreas. **B).** Representative image of DMX from retrograde-injected mouse

**Figure S5.**
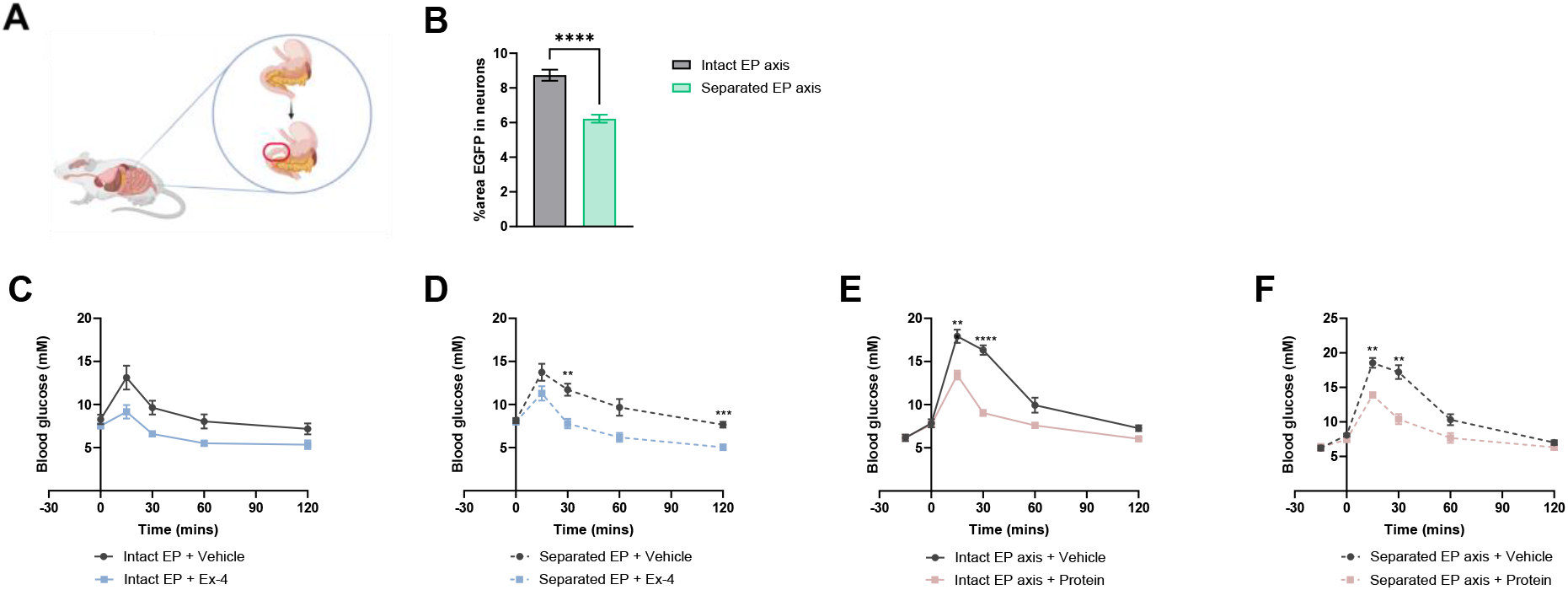
Enteropancreatic separation model validation. **A).** Schematic depicting surgical separation of proximal duodenum and pancreas in mice. **B).** Quantification of separation of the proximal duodenum and pancreas expressed as percentage eGFP expression within neurons in the LMMP of the proximal duodenum. **C).** Intraperitoneal glucose tolerance test (IPGTT) with 15-minute intraperitoneal (IP) pretreatment of Ex-4 or vehicle in mice with an intact connection between the proximal duodenum and pancreas. n=6/group. **D).** IPGTT with Ex-4 or vehicle in mice with a severed connection between the proximal duodenum and pancreas. n=7/group. **E).** IPGTT with 15-minute pretreatment of oral gavaged (OG) whey protein (10% weight/vol) or vehicle in mice with an intact connection between the proximal duodenum and pancreas. n=12/group. **F).** IPGTT with 15-minute pretreatment of OG 10% whey or vehicle in mice with a severed connection between the proximal duodenum and pancreas. n=8-10/group. Percentage area EGFP data was analysed using unpaired t-test. All IPGTT data were analysed using a 2-way ANOVA with Tukey’s post-hoc analysis and embedded AUC data were analysed using paired t-test.

**Figure S6.**
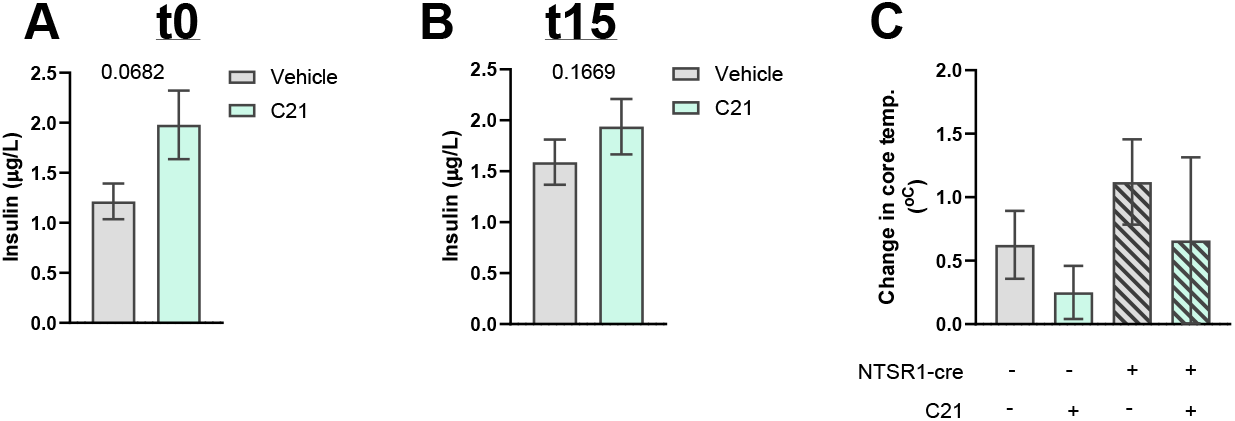
Enteropancreatic chemogenetic model validation. *NTSR1-WT* and *NTSR1-cre* mice were injected with AAVrg-hSyn-DIO-hM3D(q) into the pancreas. **A).** Plasma insulin levels measured using Mouse Insulin ELISA 15 minutes post-IP C21 or vehicle a (t0 on IPGTT). n= 13/group. **B).** Plasma insulin levels measured using Mouse Insulin ELISA 30 minutes post-IP C21 or vehicle administration and 15 minutes post-IP glucose load (t15 on IPGTT). n= 13/group. **C).** No change in core temperature following vehicle or C21 IP in *NTSR1-WT:Dq-DREADD*and *NTSR1-cre:Dq-DREADD*. n=3-4/group. Insulin data analysed using paired t-test. Change in core temperature between groups analysed using a 1-way ANOVA.

**Figure S7.**
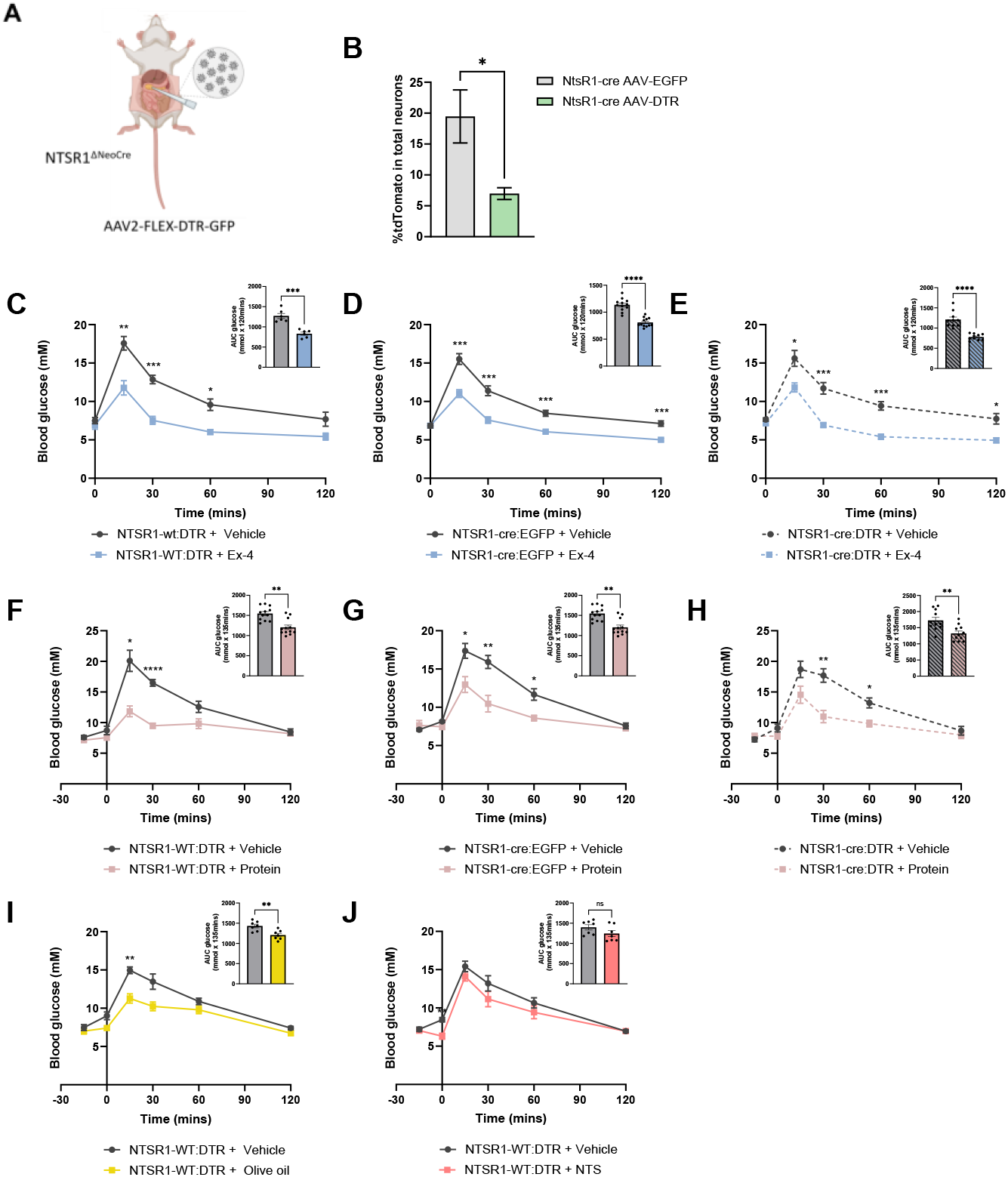
Enteropancreatic DTR model validation. **A).** Schematic depicting injection of AAV2-FLEX-DTR-GRP into the pancreas. **B).** Quantification of ablation of NTSR1-expressing neurons expressed as percentage tdTomato expression within neurons in the LMMP of the proximal. **C).** IPGTT with intraperitoneal (IP) Ex-4 or vehicle 15 minutes prior to glucose load in *NTSR1-WT* mice injected with AAV2-FLEX-DTR-GFP. n=6/group. **D).** IPGTT with IP Ex-4 or vehicle 15 minutes prior to glucose load in *NTSR1-cre* mice injected with AAV2-pCAG-FLEX-EGFP-WPRE (viral control). n=12/group. **E).** IPGTT with IP Ex-4 or vehicle 15 minutes prior to glucose load *in NTSR1-cre* mice injected with AAV2-FLEX-DTR-GFP. n=10/group. **F).** IPGTT with oral gavaged (OG) 10% whey or vehicle 15 minutes prior to glucose load in *NTSR1-WT* mice injected with AAV2-FLEX-DTR-GFP. n=7/group. **G).** IPGTT with OG 10% whey or vehicle 15 minutes prior to glucose load in *NTSR1-cre* mice injected with AAV2-pCAG-FLEX-EGFP-WPRE. n=12/group. **H).** IPGTT with OG 10% whey or vehicle 15 minutes prior to glucose load in *in NTSR1-cre* mice injected with AAV2-FLEX-DTR-GFP.

